# Climate isolation and percolation as drivers of terrestrial vertebrate richness

**DOI:** 10.64898/2026.07.07.736971

**Authors:** Marcio R. Pie

## Abstract

Climate is a strong predictor of global species richness, but the effects of climatic conditions are difficult to separate from the geography of the climates themselves. Recent work in climate space has shown that the area and isolation of discrete climatic conditions explain broad-scale richness gradients, yet the internal spatial cohesion of those climates remains poorly characterized. Here, we introduce climate percolation as a complementary descriptor of climate geography, measuring the degree to which the total area of a climate bin is concentrated within effectively connected fragments. Using global range maps for amphibians, birds, mammals and reptiles, we quantified species richness across a two-dimensional climate space defined from 12 climatic variables and evaluated the independent and joint effects of climate area, climate isolation and climate percolation across multiple climate-space resolutions. Climate isolation and percolation were strongly coupled: their first joint axis explained, on average, more than 95% of their shared variation, revealing a dominant gradient of climate fragmentation along which geographically isolated climates are also internally subdivided. Despite this collinearity, percolation consistently outperformed isolation in cross-validation across all four vertebrate groups, with particularly strong predictive gains for birds and mammals. The largest improvements, however, came from the shared isolation–percolation axis, indicating that vertebrate richness in climate space is more strongly associated with the integrated geographical structure of climates than with either inter-fragment distance or internal cohesion alone. These results suggest that climate fragmentation is a multidimensional property of environmental space, combining both the distance among climate fragments and the dominance structure of connected areas. By extending climate-space approaches from area and isolation to percolation, our framework provides a more complete description of how the geography of climate may shape global richness gradients and offers a structural basis for anticipating how future changes in climate connectivity could alter biodiversity patterns.

## Introduction

Understanding the uneven distribution of species richness across the Earth has been a central goal of biogeography and macroecology for more than a century. Climate, in particular, has long been implicated as a primary correlate of broad-scale diversity gradients, with warmer, wetter and more energy-rich regions generally supporting more species than colder or drier environments (Gaston, 2000; Hawkins et al., 2003; Currie et al., 2004; Hillebrand, 2004; Field et al., 2009). Yet this dominant explanatory tradition has an important limitation: it tends to conflate the intrinsic properties of a given climate, such as its thermal regime, seasonality and moisture balance, with the geographical circumstances under which that climate occurs on Earth, including the amount of land it occupies and the degree to which that land is spatially aggregated or fragmented. Disentangling these dimensions requires a shift from geographical space to environmental space. The conceptual basis for such a shift was provided by Hutchinson (1957), who formalized the distinction between a species’ ecological niche, represented as a region in multidimensional environmental hypervolume, and the geographical biotope that it occupies. As developed further by Colwell and Rangel (2009), Hutchinson’s duality describes a reciprocal mapping between these domains: each point in environmental space corresponds to one or more locations on the Earth’s surface, whereas each geographical location maps to a single position in environmental space. This duality makes it possible to analyze biodiversity patterns not only in conventional geographical coordinates, but also directly in climate-defined environmental space, where the unit of analysis is a discrete climatic condition rather than a grid cell on a map (Broennimann et al., 2012).

The first formal empirical test of the relationship between the structure of climatic niche space and species richness was provided by Meyer and Pie (2018). Using more than 10,000 species of amphibians and mammals, they showed that environmental prevalence, defined as the geographical availability of different climatic combinations, is a strong predictor of the number of species occupying each region of climatic niche space. Their results challenged the view that climatic conditions affect richness only through their intrinsic physiological or ecological properties, showing instead that much of the apparent climate–richness relationship reflects differences in how common those conditions are geographically. Meyer and Pie (2018) therefore operationalized Hutchinson’s duality in macroecology and demonstrated that the geography of climate matters for understanding global diversity patterns. This interpretation is consistent with broader niche theory, in which species distributions are understood as the joint outcome of environmental suitability, geographic accessibility and the mapping between environmental and geographic spaces (Soberón & Nakamura, 2009), as well as with the increasing use of multidimensional hypervolumes to quantify ecological niches and diversity patterns in environmental space (Blonder et al., 2014). This perspective was extended theoretically by Soberón (2019), who showed from first principles that species–area relationships can emerge from the structure of niche space: as the extent of environmental space available to a region increases, more species’ fundamental niches are included, thereby increasing observed richness. These theoretical expectations were tested at a global scale by Coelho et al. (2023), who evaluated both the area and spatial isolation of climate bins as predictors of species richness across all four tetrapod groups, encompassing more than 30,000 species. Their study showed that considering both climate itself and the geographical attributes of climatic conditions, specifically their total area and the mean distance among geographically disjunct fragments sharing the same climate, explains nearly 90% of global variation in vertebrate species richness. This emphasis on the geography of climatic conditions also aligns with process-based macroecological models showing that global biodiversity patterns emerge from the interaction between ecological tolerances, geographic structure, dispersal and evolutionary dynamics over deep time (Rangel et al., 2018). By shifting the unit of analysis from geographical grid cells to climate-defined bins, Coelho et al. (2023) demonstrated that the geography of climate carries explanatory power for biodiversity patterns that is largely hidden in conventional geographical-space analyses. Graham et al. (2025) further developed this environmental-space framework, showing that it can reshape the interpretation of species richness, spatial turnover, functional diversity and other macroecological patterns that have traditionally been described only in geographical coordinates.

Despite this progress, a key component of how climates are distributed across Earth remains underexplored. Climate isolation, as measured by Coelho et al. (2023), captures the between-fragment dimension of geographical structure: how far apart the spatially disjunct patches of a climate are, for example as mean pairwise distance, *D* = [2/(*n*(*n* - 1))] Σ*_i<j_ d_ij_*, where n is the number of fragments and *d_ij_* is the geographic distance between fragments *i* and *j*. Yet isolation alone is incomplete. A climate may be widely scattered while dominated by one large landmass, or have low inter-fragment distances while being divided into many similarly sized patches. These scenarios carry different implications for colonization, diversification, extinction and faunal assembly because fragmentation depends not only on area but also on subdivision, connectivity and isolation (Hanski & Ovaskainen, 2000; Haddad et al., 2015; Fahrig, 2017). We therefore introduce climate percolation as a complementary measure of within-climate structural cohesion: the extent to which the total area of a climate bin is concentrated within effectively connected fragments. High percolation indicates that most area is concentrated in one or a few large connected fragments, whereas low percolation indicates subdivision among many smaller fragments (Fig. 1).

**Figure 1.**
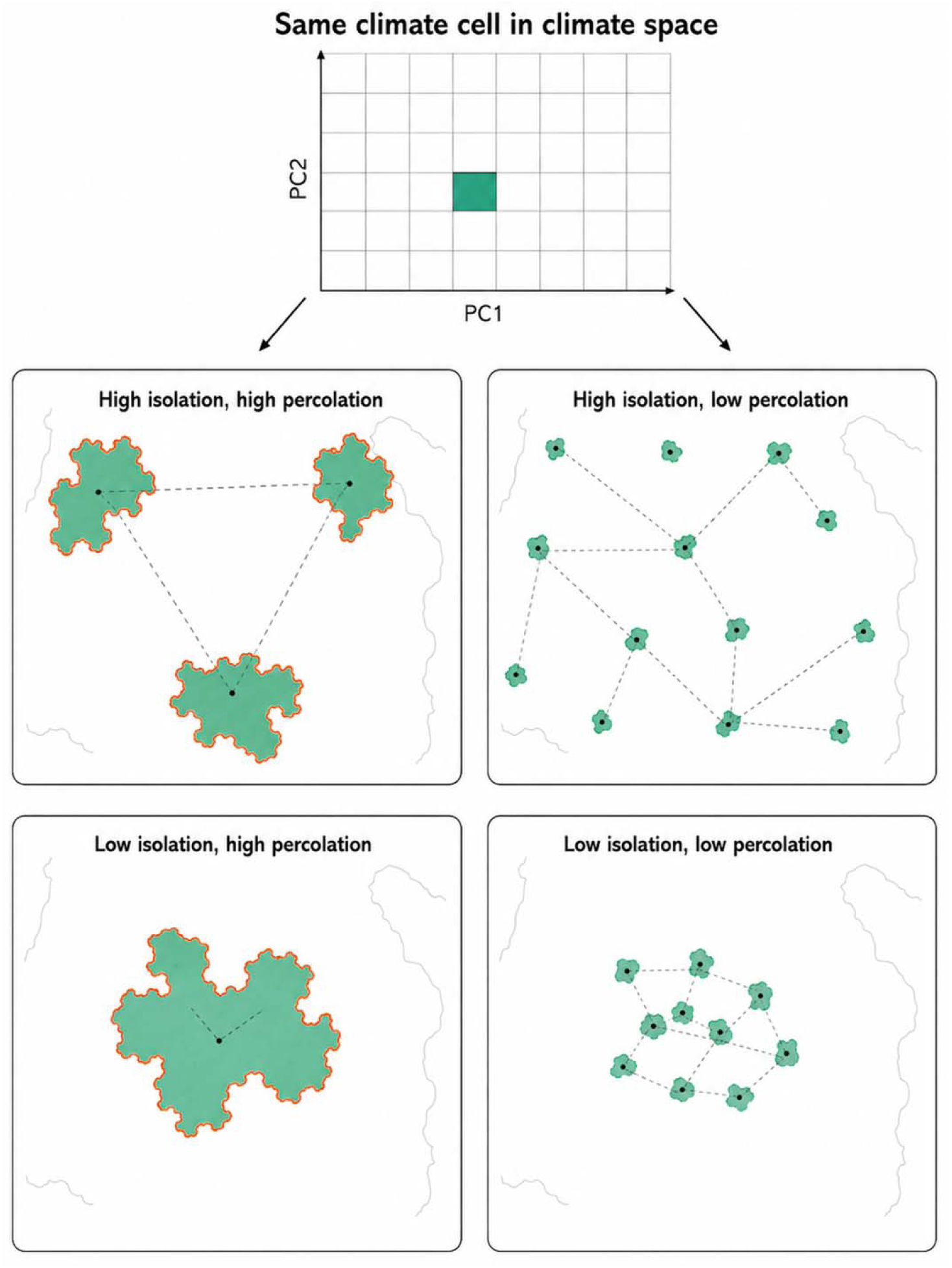
Conceptual distinction between climate isolation and climate percolation. Isolation was defined from geodesic distances among connected climate fragments, whereas percolation was defined from the connected geographic expression of a climate bin.

Following Coelho et al. (2023) and Graham et al. (2025), we adopt a climate-space framework in which each unique combination of climatic conditions constitutes a discrete environmental bin, analogous to a grid cell in geographical space. We extend the framework of Coelho et al. (2023) by decomposing climate connectivity into two complementary metrics: climate isolation, which captures the mean geodesic distance among geographically disjunct fragments of a climate bin, and climate percolation, which captures the degree to which the total area of a climate bin is concentrated in effectively connected geographical fragments. We assess the independent and joint contributions of these metrics to terrestrial vertebrate species richness and ask whether climate isolation and climate percolation carry distinct, non-redundant explanatory power for global biodiversity patterns.

## Methods

### Species richness in climate space

We evaluated the relationship between species richness and the geographical structure of climate for four terrestrial vertebrate groups: amphibians, birds, mammals and reptiles. Amphibian and mammal distributions were obtained from IUCN Red List range maps (IUCN, 2023), retaining polygons corresponding to extant, native and resident occurrences when these fields were available (presence = 1, origin = 1, seasonal = 1), while also retaining records with missing values for these attributes. Bird distributions were obtained from the BirdLife 2024.2 range maps (BirdLife International & Handbook of the Birds of the World, 2024). We used the All_Species polygon layer after exporting a filtered GeoPackage retaining current, native and resident polygons (presence = 1, origin = 1, seasonal = 1). Reptile distributions were obtained from the Global Assessment of Reptile Distributions (GARD v1.7; Roll et al., 2017; Caetano et al., 2022), which provides expert-validated range polygons for 10,914 terrestrial species.

### Climate-space construction

We defined climate space using 12 environmental variables: 11 CHELSA v2.1 bioclimatic variables for the reference period 1981–2010 (bio1, bio2, bio3, bio4, bio5, bio6, bio7, bio12, bio13, bio14 and bio15; Karger et al., 2017) and annual potential evapotranspiration (PET). PET was calculated as the sum of 12 monthly layers from the Global ET0/PET v3.1 product (Zomer et al., 2022). All climatic variables were aggregated to an equal-area Behrmann grid of approximately 110 km resolution using cell-wise means, following the approach of Coelho et al. (2023). This resolution has been identified as appropriate for global macroecological analyses using range-map data, being less prone to false species presences at large spatial extents (Hurlbert & Jetz, 2007; Belmaker & Jetz, 2011).

A principal components analysis (PCA) was performed on the correlation matrix of standardized climatic variables computed across all terrestrial grid cells with complete data. The first two principal components (PC1 and PC2), which together captured approximately 80% of global climatic variation (Coelho et al., 2023), were used to define a two-dimensional orthogonal climate space representing the principal axes of thermal and moisture variation among terrestrial environments. This space was then discretized at five resolutions by dividing each climate axis into 20, 30, 40, 50 and 60 equal intervals, following sensitivity analyses in Coelho et al. (2023) confirming result robustness across this range. Each geographical grid cell was assigned to the climate bin corresponding to its PC1 and PC2 scores.

For each climate bin, we computed three geographical descriptors used in subsequent analyses: (i) total geographical area (its environmental prevalence, sensu Meyer and Pie, 2018), measured as the total area of equal-area geographical grid cells assigned to that bin; (ii) climate-space position, represented by the within-bin arithmetic mean scores on PC1 and PC2; and (iii) the full set of geographical grid cells sharing that climatic condition, which served as the basis for the fragmentation metrics described below. Species richness in each climate bin was calculated as the number of species whose retained range intersected at least one geographical grid cell assigned to that bin.

### Climate isolation

Following Coelho et al. (2023), we characterized the between-fragment dimension of geographical structure using climate isolation. For each climate bin, we first identified all spatially connected geographical fragments using eight-neighbor adjacency among grid cells of the Behrmann raster, implemented via connected-component labelling in terra (Hijmans, 2024). The centroid of each fragment was then computed as the mean coordinate of its constituent grid cells, and the mean geodesic distance among all pairs of fragment centroids belonging to the same climate bin was taken as the isolation value for that bin. For climate bins with more than 1000 fragments, this mean distance was estimated from an area-weighted sample of up to 1000 fragments. Geodesic distances were computed using the geosphere package (Hijmans, 2022). Because the resulting isolation values were strongly right-skewed, they were log-transformed prior to analysis using the log1p transformation (i.e., log(isolation + 1)) to reduce the influence of outliers and improve model residual behavior.

### Climate percolation

We introduced climate percolation as a complementary measure of within-climate structural cohesion. For each climate bin, we identified all connected geographical fragments and calculated the fraction of total climate-bin area represented by each fragment, 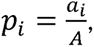 where a is the area of fragment *i* and *A* is the total area of that climate bin. Climate percolation was then measured as 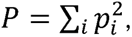 which is equivalent to the probability that two geographical grid cells drawn at random from the same climate bin belong to the same connected fragment, and is formally related to the degree-of-landscape-division and effective mesh-size statistics introduced by Jaeger (2000). Values of P approaching one indicate that nearly all of a climate bin’s geographical area is concentrated within a single dominant, continuously connected fragment. Lower values indicate that the same total area is distributed among multiple fragments of comparable size. Unlike metrics based solely on the largest connected component, P is sensitive to the full distribution of fragment sizes and therefore provides a richer characterization of spatial subdivision. Climate percolation was used on its natural scale (0, 1] in all analyses.

### Residualization and isolation–percolation decomposition

Because climate isolation and climate percolation are geometrically related (both being determined by the same underlying spatial arrangement of fragments) their raw values are expected to covary. To evaluate independent contributions, we used two complementary approaches. First, we residualized each metric against log-transformed climate area and climate-space position (PC1 and PC2) using Gaussian GAMs with shrinkage smooths fitted via restricted maximum likelihood (REML; Wood, 2011). The resulting residuals capture variation in isolation or percolation that is not attributable to the size or climatic identity of the bin, hereafter termed area- and position-adjusted residuals, respectively. We further computed unique residuals for each metric by residualizing isolation against area, position and percolation, and percolation against area, position and isolation, so as to isolate the contribution of each metric after explicitly accounting for the other. Second, to characterize the dominant axis of covariation between isolation and percolation and any orthogonal contrast between them, we performed a two-variable PCA on standardized log-transformed isolation and untransformed percolation values separately for each climate-space resolution. The first axis was oriented so that increasing scores correspond to increasing isolation and decreasing percolation — a gradient we term the shared climate-fragmentation axis — and the second axis captured the orthogonal, independent contrast between the two metrics.

### Statistical models

All primary richness models were negative-binomial GAMs fitted with the mgcv v. 1.9-1 package (Wood, 2011; Wood et al., 2016; Wood, 2017) using REML smoothness selection, shrinkage smooths and smooth-term selection. The negative-binomial family was chosen over the Poisson family to account for overdispersion in species richness counts, which is common in climate-space analyses where richness varies by several orders of magnitude across bins. For each taxonomic group and climate-space resolution, we first fitted a baseline model containing smooth terms for log-transformed climate area, PC1 and PC2. We then compared this baseline against a series of augmented models that added, in turn: (i) the area- and position-adjusted residual of isolation alone; (ii) the area- and position-adjusted residual of percolation alone; (iii) both residual metrics simultaneously; (iv) the unique residuals of isolation and percolation; and (v) the first and second PCA axes of the joint isolation–percolation structure. All smooth terms were constrained to a maximum basis dimension of k = 10, reduced when necessary to remain consistent with sample size. Model support was quantified as the gain in proportion of null deviance explained relative to the baseline model, and as the reduction in AIC (ΔAIC).

Predictive performance was assessed via five-fold cross-validation repeated five times independently for each taxon–resolution combination, yielding 25 held-out evaluation sets per model configuration. In each replicate, models were refitted on four folds and their predictions evaluated on the remaining fold using Poisson deviance as a measure of predictive error for count data. Predictive gain was defined as the mean reduction in out-of-sample deviance relative to the baseline model, averaged across all 25 replicates. To make these results as comparable to those of Coelho et al. (2023), we also replicated their modelling framework using Poisson GAMs with a basis dimension of k = 4 and the default mgcv fitting criterion. These models used the same richness, isolation and percolation data but adopted the Poisson family and a more constrained smooth basis, providing a direct point of comparison with the published framework. Because the Poisson family does not explicitly model overdispersion, these supplementary models may underestimate predictive uncertainty relative to the primary negative-binomial models; we therefore treat their results as corroborating evidence rather than primary inference.

All analyses were conducted in R version 4.2.1 (R Core Team, 2024), with spatial operations performed using terra v.1.7-65 (Hijmans, 2024) and sf v1.1-0 (Pebesma, 2018), geodesic distance calculations using geosphere v. 1.5-18 (Hijmans, 2022), data handling using data.table v.1.14.10 (Barrett et al. 2023), visualization using ggplot2 v.4.0.2 (Wickham et al. 2026), patchwork v.1.3.0 (Pedersen 2024) and scales v.1.4.0 (Wickham et al. 2025), and statistical modelling using mgcv v1.9-1 (Wood, 2011; Wood et al., 2016; Wood, 2017).

## Results

Climate isolation and effective connected fraction were strongly coupled across climate-space bins, with more isolated climates tending to have lower internal connectivity (Fig. 2). Using effective connected fraction as the percolation metric, the first axis of the two-variable PCA explained, on average, 95.4% of the joint variation in isolation and percolation across climate-space resolutions, whereas the second axis explained 4.6% (Fig. 2). The dominant axis therefore represented a shared gradient of climate fragmentation, along which more isolated climates were also less geographically connected.

**Figure 2.**
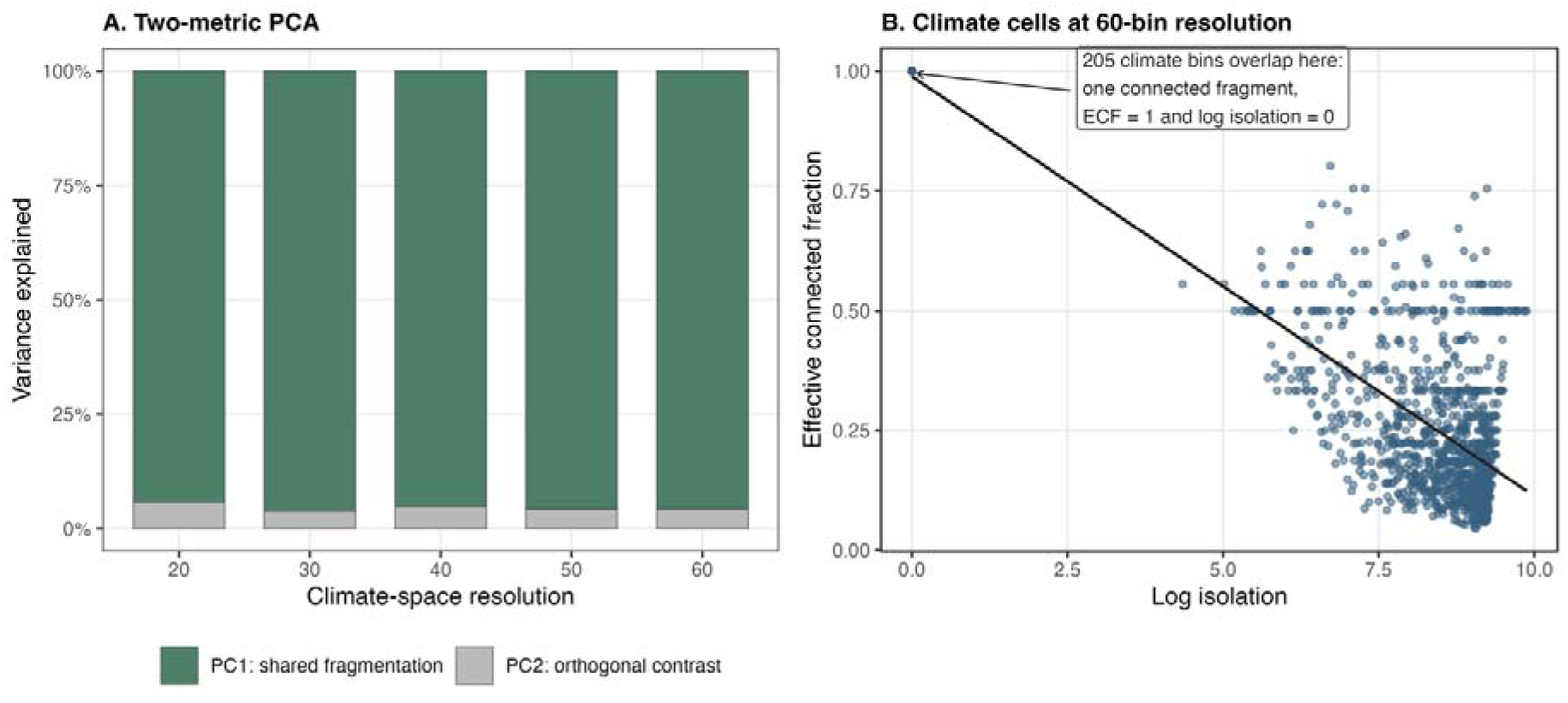
Coupling between climate isolation and effective connected fraction. (A) Proportion of variance explained by the first two axes of a two-variable PCA performed on standardized log-transformed climate isolation and effective connected fraction, separately for each climate-space resolution. PC1 represents the shared fragmentation axis, contrasting high isolation with low internal connectivity, whereas PC2 represents the residual orthogonal contrast between the two metrics. (B) Empirical relationship between log isolation and effective connected fraction across climate-space bins at the finest resolution, 60 bins per climate axis. Each point represents one occupied climate bin; the fitted line shows the overall linear trend. The annotation indicates that 205 climate bins overlap at the upper-left point, where effective connected fraction equals 1 and log isolation equals 0. These bins correspond to climates represented by a single connected geographic fragment, for which among-fragment distance is zero and all occupied area belongs to the same fragment.

Cross-validation provided stronger support for percolation (Fig. 3; Table 1). Mean predictive improvement for residual percolation exceeded that for residual isolation in all four groups: amphibians (148.6 vs. 110.3 deviance units), birds (742.3 vs. 517.5), mammals (277.4 vs. 67.6) and reptiles (395.1 vs. 192.7). Residual percolation outperformed residual isolation in 4 of five resolutions for amphibians, 5 of five for birds, 5 of five for mammals and 3 of five for reptiles (Fig. 4; Table S1). Reptiles showed the weakest resolution-level consistency, but residual percolation still outperformed residual isolation in 3 of five resolutions and had the larger mean predictive gain.

**Figure 3.**
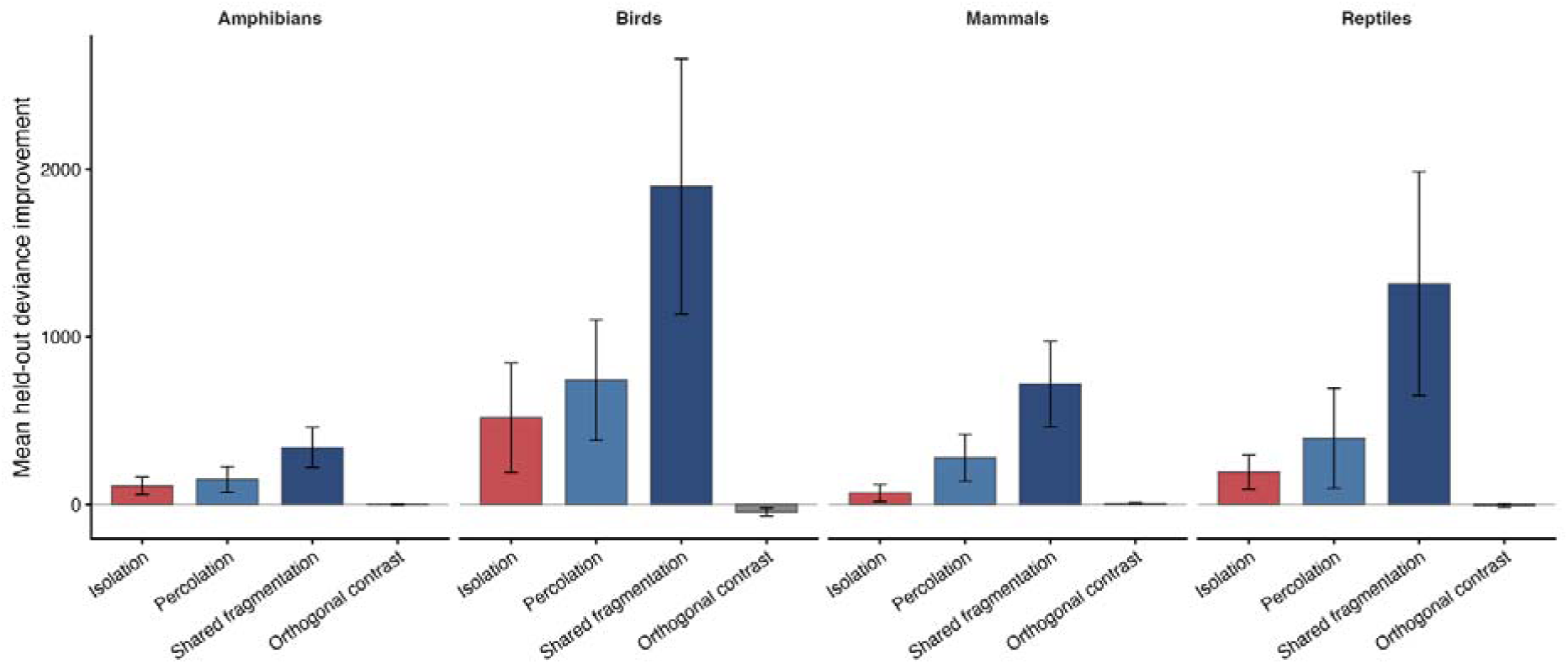
Cross-validated predictive support for isolation, percolation and their shared fragmentation structure. Bars show mean improvement in held-out Poisson deviance relative to the baseline model containing environmental prevalence and climate-space position; error bars are standard errors across climate-space resolutions.

**Figure 4.**
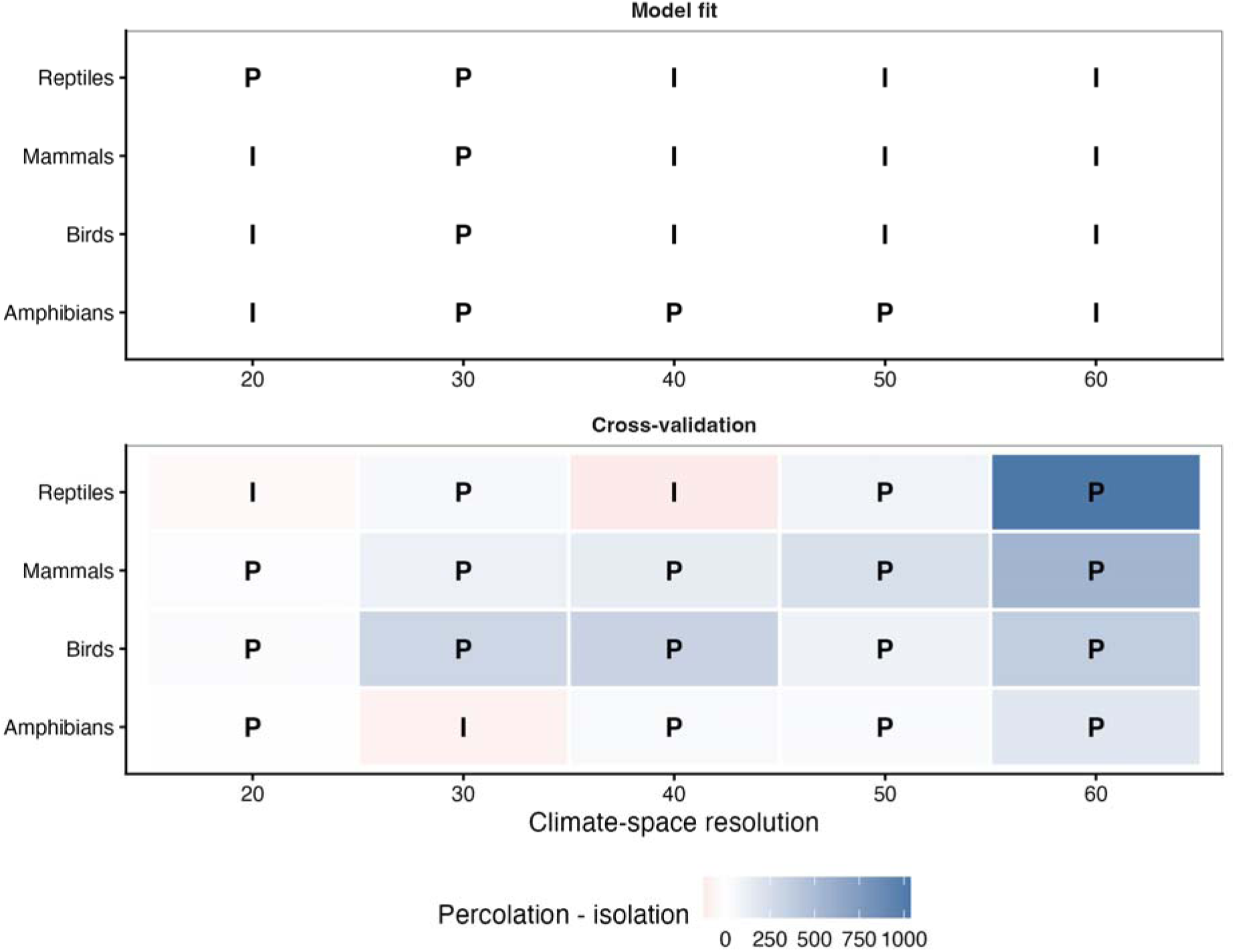
Relative support for percolation and isolation across taxa and climate-space resolutions. Colors show percolation minus isolation, with positive values favoring percolation and negative values favoring isolation. Letters indicate the stronger metric in each taxon-resolution combination.

**Table 1.** Summary of predictive support by taxonomic group. Values are mean held-out deviance improvements across climate-space resolutions, the number of resolutions in which each residual metric outperformed the other, and the mean variance explained by the shared fragmentation axis. CV (cross-validation support) gains are mean reductions in held-out Poisson deviance relative to the baseline model across climate-space resolutions.

| Taxon | CV gain: | CV gain: | CV gain: shared | CV gain: |  | Isolation |
| --- | --- | --- | --- | --- | --- | --- |
|  | isolation | percolation | fragmentation | orthogonal contrast | Percolation stronger (n/5) | stronger (n/5) |
| Amphibians | 110.3 | 148.6 | 340.5 | -0.1 | 4 | 1 |
| Birds | 517.5 | 742.3 | 1,898.8 | -45.0 | 5 | 0 |
| Mammals | 67.6 | 277.4 | 717.9 | 6.4 | 5 | 0 |
| Reptiles | 192.7 | 395.1 | 1,316.6 | -6.1 | 3 | 2 |

In the main negative-binomial GAMs, the unique effects of isolation and percolation were modest and varied among taxa and climate-space resolutions (Fig. 4; Table S1). For amphibians, residual percolation produced larger in-sample gains than residual isolation in 3 of five resolutions. For birds and mammals, residual isolation produced larger gains in 4 of five resolutions, and for reptiles it was stronger in 3 of five resolutions. Mean gains in deviance explained were small across all groups, indicating limited support for strong independent in-sample effects of either metric after accounting for environmental prevalence and climate-space position.

The strongest and most consistent signal was associated with the shared isolation-percolation fragmentation axis (Fig. 3; Table 1). In cross-validation, this axis produced the largest mean predictive gains for all taxonomic groups: 340.5 deviance units for amphibians, 1898.8 for birds, 717.9 for mammals and 1316.6 for reptiles. These improvements were substantially larger than those obtained from residual isolation or residual percolation alone, indicating that the integrated geographic structure of climates was more predictive of richness than either component considered separately. The sensitivity analyses using Poisson GAMs, as used by Coelho et al. (2023), gave even stronger relative support to percolation (Table S2). In these models, residual percolation produced larger gains than residual isolation in 3 of five resolutions for amphibians, 2 of five for birds, 5 of five for mammals and 3 of five for reptiles. Mean gains for residual percolation were also higher than those for residual isolation in all four taxonomic groups.

Overall, the results indicate that climate percolation, when measured as effective connected fraction, has greater relative predictive support than isolation in several comparisons, particularly in cross-validation and for mammals. However, the main conclusion is that isolation and percolation primarily describe a common axis of climate fragmentation. Terrestrial vertebrate richness in climate space is therefore more strongly associated with the combined geographic structure of climates, high isolation coupled with low connectivity, than with independent effects of isolation or percolation alone.

## Discussion

We extended the climate-space framework of Coelho et al. (2023) by introducing climate percolation as a complementary descriptor of how the total geographical area of a climate bin is distributed among its spatially connected fragments, and by asking whether this metric carries independent or joint predictive power for terrestrial vertebrate species richness over and above climate isolation. Three main conclusions emerge. First, climate isolation and climate percolation are strongly collinear across climate bins, with over 95% of their joint variation shared along a single dominant axis of climate fragmentation. Second, despite this collinearity, percolation consistently outperforms isolation in out-of-sample cross-validation across all four taxonomic groups, with the largest relative advantage over isolation in mammals. Third, the shared fragmentation axis, capturing the combined signal of widely separated and internally subdivided climates, is by far the strongest structural predictor of species richness in climate space, yielding predictive gains larger than those attributable to either metric individually.

The most striking result is the near-complete coupling between climate isolation and percolation. That two metrics conceived and measured from distinct geometrical properties — inter-fragment distances versus intra-bin area distribution — should share more than 95% of their joint variation implies a pervasive empirical regularity in the global geography of climate. Climates whose geographical footprint is distributed among many similarly sized fragments tend also to have those fragments scattered far apart across the Earth’s surface, whereas climates dominated by a single large continuous landmass tend to have their remaining fragments in closer geographic proximity. This regularity likely reflects shared forcing by the same deep-time geological and climatic processes, e.g. continental drift, mountain orogeny, glacial–interglacial cycles, that simultaneously determine both how far climate patches are scattered and how evenly the climate’s total area is distributed among them (Hagen et al., 2021; Rangel et al., 2018).

The dominance of this shared axis has an important interpretive consequence: it makes isolation and percolation largely redundant as proxies for the same underlying gradient of climate spatial fragmentation, at least at the resolution and spatial extent studied here. Although percolation showed stronger predictive support in cross-validation, neither metric alone captured the dominant structural signal as directly as the shared axis. The most informative unit of macroecological inference is therefore the fragmentation axis itself, a gradient that describes whether a climate’s geographical distribution is cohesive or diffuse, rather than either of its two components. This conclusion parallels observations in landscape ecology that single-metric characterizations of habitat spatial structure are rarely sufficient: the consequences of fragmentation for biodiversity depend on the joint configuration of patch sizes, patch isolation and overall spatial coherence in ways that no single distance- or area-based metric can fully capture (Gustafson, 1998; Fahrig, 2003; Fahrig, 2017; Haddad et al., 2015).

Despite their tight empirical coupling, climate percolation consistently outperforms climate isolation in predictive cross-validation, with the largest relative advantage over isolation in mammals. This asymmetry suggests that the two metrics may emphasize different components of the mechanisms by which climate fragmentation affects diversity. Climate isolation captures a long-range, inter-fragment dimension: how far the spatially disjunct patches of a climate bin are from one another. As a metric of long-range connectivity, it is most relevant for processes that operate over geological timescales and continental distances, including the interruption of gene flow across major biogeographic barriers and the independent evolutionary differentiation of regional biotas that underpins the area-isolation hypothesis (Coelho et al., 2023). Climate percolation, by contrast, captures an intra-bin, structural dimension: whether the total area of a climate bin is concentrated in a single dominant, continuously connected fragment or dispersed among many smaller patches of roughly comparable size. A climate bin with low percolation has its area subdivided among multiple fragments, reducing the dominance of any single contiguous arena in which populations and lineages might persist and accumulate. The ecological implications may extend from population dynamics, where smaller fragments tend to support smaller populations with higher extinction probabilities, to macroevolutionary dynamics: fragmentation into multiple smaller, more evenly sized patches may reduce the likelihood that any one fragment functions as a stable refugium or evolutionary cradle over the temporal scales relevant to speciation and clade diversification (Hanski & Ovaskainen, 2000; Saupe et al., 2019; Hagen et al., 2021).

The taxonomic pattern suggests that the relative importance of percolation versus isolation depends on the scale at which dispersal limitation, climatic tracking and range subdivision interact. Mammals showed the largest relative advantage of percolation over isolation, whereas birds showed the largest absolute predictive gains for both percolation and the shared fragmentation axis. This contrast is biologically plausible: birds generally have high dispersal capacity, so distances among climate fragments may be less limiting than the extent to which suitable climates form large, internally cohesive arenas. Mammals, by contrast, may be more sensitive to the internal dominance structure of climate distributions when climatic niche conservatism and slower niche evolution make fragmented climate space difficult to reintegrate over deep time. Process-based simulations of global vertebrate diversity support the broader plausibility of this mechanism, showing that tropical richness gradients can emerge from higher rates of range disjunction driven by spatio-temporal climatic change, especially precipitation, and that niche conservatism can amplify diversification by limiting adaptive tracking of shifting climate conditions (Saupe et al., 2019; Qiao et al., 2024). The weaker resolution-level consistency in reptiles may reflect the heterogeneity of this group, which spans wide variation in dispersal ability, thermal physiology and habitat specificity, so the relative importance of isolation and percolation may depend strongly on lineage-specific life history and physiology (Brodie & Mannion, 2023).

Our results contribute to the long-standing debate over why species richness is concentrated in the tropics (Mittelbach et al., 2007; Brodie & Mannion, 2023). The shared climate-fragmentation axis we identify is not randomly distributed across the globe: Coelho et al. (2023) showed that tropical climates occupy larger total geographic areas and are more fragmented than polar climates, contradicting the view that the tropics are simply more environmentally common. The present results extend this picture by demonstrating that the percolation dimension of fragmentation, how internally subdivided a climate is, mirrors the isolation dimension, so that tropical climates are characterized by both large inter-fragment distances and low within-bin spatial cohesion. This means that both large-scale geographical properties examined here, mean fragment distance and area dominance structure, point toward the same empirical conclusion: the fragmented, multi-cradle structure of tropical climate conditions is a major geographic correlate of the greater vertebrate species richness observed there, consistent with evolutionary perspectives that emphasize the tropics as both sources and reservoirs of diversity (Jablonski et al., 2006; Mittelbach et al., 2007).

This interpretation is mechanistically grounded in process-based macroecological models. Saupe et al. (2019) demonstrated, using eco-evolutionary simulations, that spatio-temporal changes in low-latitude precipitation drive range disjunctions among populations that share the same climatic niche, thereby generating high rates of allopatric speciation that strongly predict the latitudinal diversity gradient. Hagen et al. (2021) showed that variation in Earth history — mountain uplift and climate change over tens of millions of years — explains the uneven distribution of diversity among tropical moist forests across continents, with historically more geologically active and climatically dynamic regions harboring greater richness through higher rates of fragmentation-induced speciation. Together, these studies suggest that the climate-space signal we document statically in present-day geography reflects a geological legacy of accumulated speciation opportunities, consistent with the broader expectation that recurrent climatic change reshapes species’ geographical distributions, alternately fragmenting and reconnecting populations through time (Dynesius & Jansson, 2000; Jansson & Dynesius, 2002). This interpretation also aligns with Saupe’s (2023) argument that any explanation for the latitudinal diversity gradient must ultimately invoke differential rates of speciation, extinction or dispersal, rather than static correlates alone. Our shared fragmentation axis provides a tractable, empirically accessible proxy for the structural conditions that modulate these rates over deep time.

Several limitations circumscribe these results. First, richness estimates derive from expert range maps, which are appropriate for global comparisons but generalize local occupancy, seasonal movement and fine-scale habitat use. This is especially relevant for birds, whose high vagility and migratory behaviour complicate range boundaries, so richness should be interpreted as broad-scale climatic association rather than local community composition. Second, both fragmentation metrics are calculated from present-day climate, a snapshot of a system that has changed substantially over geological and Quaternary timescales. Paleoclimatic sequences of climate-bin expansion, contraction and coalescence are likely more directly relevant to current diversity patterns (Rangel et al., 2018; Saupe et al., 2019; Svenning et al., 2015), making time series of percolation and isolation a natural extension. Third, the 110 km grain is appropriate for global range-map analyses (Hurlbert & Jetz, 2007) but cannot resolve fine-scale climatic heterogeneity in mountains, which often retain positive residual richness after climate geography is considered (Coelho et al., 2023; Rahbek et al., 2019). Finally, we analyse species richness only. Whether percolation complements isolation for turnover, functional diversity and phylogenetic dispersion remains an open question.

These results also have implications for anticipating biodiversity change. Most climate-change assessments emphasize shifts in range edges and the velocity at which isotherms move across landscapes (Loarie et al., 2009; Brito-Morales et al., 2018), implicitly assuming that species can track preferred climates when movement rates are tractable. Our results suggest an additional risk: even if the total area of a climate bin persists, declining percolation could divide one dominant fragment into several smaller patches, reducing the size and continuity of the arenas in which populations persist. Conversely, future coalescence of fragments could increase long-term climatic cohesion, although too slowly to matter for decadal conservation planning. Incorporating percolation with area and isolation into warming projections would therefore provide a more structurally complete view of climatic risks to vertebrate diversity, complementing approaches based on climate connectivity and dispersal through fragmented landscapes (McGuire et al., 2016).

## Acknowledgments

We thank F. S. Caron for valuable comments on the manuscript. M.R.P. was funded by a grant from Conselho Nacional de Desenvolvimento Científico e Tecnológico (CNPq), Brazil (303491/2024).

## Data Accessibility

The species range data used in this study were obtained from publicly available sources: IUCN Red List range maps for amphibians and mammals, BirdLife International and Handbook of the Birds of the World range maps for birds, and the Global Assessment of Reptile Distributions database for reptiles. Climate data were obtained from CHELSA v2.1 and the Global ET0/PET v3.1 product.

## Code Availability

All R scripts used to construct climate space, calculate climate isolation and climate percolation, fit statistical models, conduct cross-validation and generate figures are deposited in Zenodo (https://zenodo.org/records/20645401?token=eyJhbGciOiJIUzUxMiJ9.eyJpZCI6ImJlZDczNzU5LWE2ZmYtNDg2ZC05NzE5LTRiYzYwM2JkODc1YSIsImRhdGEiOnt9LCJyYW5kb20iOiJjZDY2MjM0MjhiZDc1N2UxY2EzMjU1NGE4YzE5ODliZiJ9.zc0BRNfMxT31IpzuP3_TX_V1r0MM6m8NwoqMoCQXzsEW0ZQpPcu2ZdE3rbtqyX4ShLeuFok9HQMYFYyosrX9YQ).

## Author Contributions

M.R.P.: Conceptualization; Methodology; Software; Formal analysis; Investigation; Data curation; Visualization; Writing - original draft; Writing - review & editing; Funding acquisition.

---

**Table S1.** Resolution-level results from the main negative-binomial GAMs and cross-validation analyses, including gains in deviance explained, held-out deviance improvements and the stronger residual metric for each taxon-resolution combination.

| Taxon | Resolution | In-sample gain: isolation | In-sample gain: percolation | In-sample gain: shared fragmentation | In-sample gain: orthogonal contrast | In-sample winner | CV gain: isolation | CV gain: percolation | CV gain: shared fragmentation | CV gain: orthogonal contrast | CV winner |
| --- | --- | --- | --- | --- | --- | --- | --- | --- | --- | --- | --- |
| Amphibians | 20 | 0.0000 | 0.0000 | 0.0000 | 0.0000 | Isolation | -9.1 | -1.4 | -5.8 | -0.1 | Percolation |
| Amphibians | 30 | -0.0082 | 0.0011 | -0.0052 | -0.0001 | Percolation | 175.2 | 95.9 | 370.9 | 6.2 | Isolation |
| Amphibians | 40 | -0.0020 | -0.0006 | -0.0030 | -0.0000 | Percolation | -7.7 | 33.8 | 213.2 | -0.5 | Percolation |
| Amphibians | 50 | -0.0033 | -0.0016 | -0.0053 | 0.0000 | Percolation | 144.1 | 181.4 | 413.0 | -6.9 | Percolation |
| Amphibians | 60 | 0.0002 | -0.0028 | -0.0055 | 0.0000 | Isolation | 249.0 | 433.4 | 711.5 | 0.8 | Percolation |
| Birds | 20 | 0.0077 | 0.0000 | 0.0024 | 0.0021 | Isolation | -41.3 | -15.8 | -45.6 | -65.1 | Percolation |
| Birds | 30 | 0.0038 | 0.0171 | 0.0077 | 0.0000 | Percolation | 376.3 | 679.5 | 1,450.9 | -26.9 | Percolation |
| Birds | 40 | 0.0118 | 0.0036 | 0.0030 | 0.0000 | Isolation | -119.0 | 206.9 | 1,143.4 | -1.0 | Percolation |
| Birds | 50 | 0.0085 | 0.0013 | 0.0019 | 0.0003 | Isolation | 679.6 | 780.9 | 2,449.6 | -132.0 | Percolation |
| Birds | 60 | 0.0087 | 0.0047 | 0.0039 | 0.0000 | Isolation | 1,692.1 | 2,060.1 | 4,495.7 | -0.2 | Percolation |
| Mammals | 20 | 0.0063 | 0.0004 | 0.0053 | 0.0000 | Isolation | -21.1 | -5.4 | 44.7 | -0.2 | Percolation |
| Mammals | 30 | 0.0001 | 0.0013 | -0.0016 | 0.0000 | Percolation | 70.8 | 178.7 | 424.2 | 1.2 | Percolation |
| Mammals | 40 | 0.0020 | 0.0013 | -0.0058 | 0.0001 | Isolation | -47.0 | 93.2 | 589.3 | 21.4 | Percolation |
| Mammals | 50 | -0.0006 | -0.0016 | -0.0113 | 0.0000 | Isolation | 93.7 | 328.9 | 977.9 | 6.7 | Percolation |
| Mammals | 60 | -0.0015 | -0.0043 | -0.0141 | 0.0000 | Isolation | 241.7 | 791.7 | 1,553.1 | 3.0 | Percolation |
| Reptiles | 20 | 0.0000 | 0.0000 | 0.0000 | 0.0000 | Percolation | 24.0 | -15.7 | 134.8 | -24.6 | Isolation |
| Reptiles | 30 | 0.0000 | 0.0000 | -0.0001 | 0.0000 | Percolation | -27.7 | 19.8 | -30.8 | -0.3 | Percolation |
| Reptiles | 40 | 0.0019 | 0.0000 | -0.0045 | -0.0000 | Isolation | 123.7 | 2.3 | 1,002.9 | 7.2 | Isolation |
| Reptiles | 50 | -0.0008 | -0.0021 | -0.0085 | 0.0000 | Isolation | 341.5 | 430.7 | 1,855.1 | -33.1 | Percolation |
| Reptiles | 60 | -0.0010 | -0.0049 | -0.0131 | 0.0000 | Isolation | 501.8 | 1,538.6 | 3,621.1 | 20.1 | Percolation |

**Table S2.** Supplementary sensitivity analysis based on the approach of Coelho et al. (2023) using Poisson GAMs with k = 4, showing residual isolation and residual percolation gains for each taxon-resolution combination. Coelho-style gains are gains in deviance explained relative to the Coelho-style baseline Poisson GAM. Delta AIC values are baseline AIC minus augmented-model AIC; positive values favor the augmented model.

| **Taxon** | **Resolution** | **Coelho-style gain:**<br>**isolation** | **Coelho-style gain:**<br>**percolation** | **Winner** | **Delta AIC:**<br>**isolation** | **Delta AIC:**<br>**percolation** |
| --- | --- | --- | --- | --- | --- | --- |
| Amphibians | 20 | 0.0022 | 0.0106 | Percolation | 982.8 | 1,581.9 |
| Amphibians | 30 | 0.0055 | 0.0122 | Percolation | 1,883.5 | 2,400.8 |
| Amphibians | 40 | 0.0042 | 0.0045 | Percolation | 1,459.5 | 1,879.2 |
| Amphibians | 50 | 0.0084 | 0.0070 | Isolation | 2,815.0 | 2,745.8 |
| Amphibians | 60 | 0.0131 | 0.0114 | Isolation | 3,701.3 | 3,775.4 |
| Birds | 20 | 0.0041 | 0.0085 | Percolation | 5,671.7 | 6,548.5 |
| Birds | 30 | 0.0104 | 0.0141 | Percolation | 12,698.5 | 12,904.4 |
| Birds | 40 | 0.0072 | 0.0072 | Isolation | 10,951.5 | 12,093.4 |
| Birds | 50 | 0.0142 | 0.0091 | Isolation | 21,733.0 | 17,528.3 |
| Birds | 60 | 0.0184 | 0.0155 | Isolation | 28,729.2 | 25,533.6 |
| Mammals | 20 | 0.0032 | 0.0087 | Percolation | 1,579.7 | 2,204.1 |
| Mammals | 30 | 0.0089 | 0.0141 | Percolation | 3,886.3 | 5,311.6 |
| Mammals | 40 | 0.0050 | 0.0075 | Percolation | 3,274.4 | 4,828.3 |
| Mammals | 50 | 0.0093 | 0.0115 | Percolation | 6,951.7 | 7,921.8 |
| Mammals | 60 | 0.0169 | 0.0193 | Percolation | 8,828.9 | 10,358.1 |
| Reptiles | 20 | 0.0021 | 0.0141 | Percolation | 1,745.0 | 3,723.6 |
| Reptiles | 30 | 0.0060 | 0.0114 | Percolation | 4,323.9 | 5,721.5 |
| Reptiles | 40 | 0.0052 | 0.0058 | Percolation | 4,665.9 | 5,112.2 |
| Reptiles | 50 | 0.0121 | 0.0083 | Isolation | 9,337.8 | 7,627.9 |
| Reptiles | 60 | 0.0167 | 0.0146 | Isolation | 11,884.3 | 10,570.7 |

